# Deep CyTOF immune profiles of ex vivo granulomatous responses to *Mycobacterium tuberculosis* differ with European and African geographical origin

**DOI:** 10.64898/2026.11.09.764574

**Authors:** Sarah Schmidiger, Erin F. McCaffrey, Jan M. Schmidt, Laura Zaragoza-Infante, Amanda Ross, Max Mpina, Anneth Tumbo, Elirehema Mfinanga, Frederick Haraka, Jerry Hella, José M. Carballido, Klaus Reither, Ainhoa Arbués, Damien Portevin

## Abstract

Genetic and environmental factors influence human immunity and modulate responses to infection. Accounting for immune heterogeneity across human populations is critical to the design of novel vaccines and host-directed interventions that would optimally benefit populations experiencing the highest disease burden. Tuberculosis (TB) burden is unevenly distributed across geographical regions and our understanding of the differential immune response to *Mycobacterium tuberculosis* (*Mtb*) across human populations remains limited. Here, we sought to evaluate the impact of geographical origin on ex vivo granulomatous responses to *Mtb* infection. We found that peripheral blood mononuclear cells (PBMCs) from healthy African donors exhibited greater control of *Mtb* growth ex vivo than those isolated from European individuals. High-dimensional single-cell profiling of the granulomatous responses by cytometry by time-of-flight (CyTOF) revealed divergent profiles based on the specimens’ geographical origin. Monocytes from European donors displayed a pro-inflammatory phenotype – correlating with *Mtb* growth –, whereas PBMCs of African individuals were enriched in activated, cytotoxic T cells – correlating with *Mtb* control. Reduced inflammatory response in African monocytes following various microbial stimulations was confirmed. Taken together, our comprehensive immune profiling indicates that environmental and genetic factors shape a cytotoxic-regulatory response that promotes bacterial control. Against the current “one-treatment-fits-all” approach, our findings support the use of specimens from intervention-relevant settings to inform the targeted design of TB vaccines and host-directed interventions.

**SIGNIFICANCE STATEMENT:** Human immunity and responses to infection vary between and within populations. Considering this immunological diversity is particularly important to the development of innovative interventions for globally unevenly distributed infectious diseases, such as tuberculosis. We employed a high-dimensional single-cell approach to characterize the divergence of immune responses to mycobacterial infection ex vivo between African and European blood specimens. The increased bacterial growth control of African specimens was associated with highly cytotoxic, partially regulatory T cells and monocyte tolerance to microbial stimulation – a protective signature, independent of mycobacteria-specific adaptive immunity, epigenetically shaped by environmental triggers and heritable genetic traits. These data suggest protective immune traits to be context-dependent and emphasize the need to conduct and validate research where interventions are needed most.

## INTRODUCTION

Influenced by intrinsic (e.g. age, sex, genetics) and extrinsic (e.g. environmental, behavioral, nutritional) factors, human immunity varies across individuals and over time.^1–3^ Pathogen exposure and vaccination elicit adaptive immune responses that persist as immune memory, enabling a faster and highly specific response upon re-exposure. Following an epigenetically mediated process termed “trained innate immunity”, innate immune cells can also become more responsive to subsequent encounters with microbes.^4^ However, chronic exposure to a pathogenic stimulus can result in tolerance of innate cells that hence become less responsive or anergic to secondary stimulation.^5,6^ Unlike adaptive immunity, innate immune training displays high levels of cross-reactivity. For example, trained immunity induced by BCG vaccination^7^ confers protection against viral and parasitic infections.^8–10^ Furthermore, immune responses to bacterial and viral pathogens vary across populations in relation to their degree of African versus European ancestry.^11–15^ To conceive innovative immunotherapeutic approaches for the management of infectious diseases, it thus appears critical to consider the immunological variance within and across human populations worldwide.

Exposure to *Mycobacterium tuberculosis* (*Mtb*), the causative agent of human tuberculosis (TB), only results in active disease in around 5% of cases.^16^ Still, this ancient pathogen causes an estimated 10 million incident cases every year and claimed the lives of 1.23 million individuals in 2024 alone.^17^ TB pathogenesis is tightly interlinked with human immunology, for immune responses can both confer protection and exacerbate disease.^18^ Therefore, human immunological diversity cannot be overlooked in developing interventions for TB. Yet, human TB immunology research is too often conducted on human samples collected in high-income, low-TB-endemicity settings, where interventions are less needed, or in clinical TB cohorts consisting of individuals with a mix of different disease presentations (e.g. latent vs active TB). In contrast, cross-contextual studies on immune responses to *Mtb* in healthy individuals that would allow characterization of protective traits are scarce. Using a clinically relevant 3D granuloma model^19^ and a 43-parameter mass cytometry-based single-cell approach (CyTOF)^20^, we compared ex vivo immune responses to *Mtb* infection of healthy individuals (n = 24) across two different geographical settings. We found African peripheral blood mononuclear cells (PBMCs) to control *Mtb* growth ex vivo significantly better than European PBMCs did, independently of *Mtb*-specific T cell memory. Functional single-cell granuloma profiles differed most strikingly according to their geographical origin. Specifically, European profiles showed pronounced pro-inflammatory monocyte responses, while African profiles were enriched in cytotoxic and regulatory CD8 T cells, which correlated with bacterial growth promotion and control, respectively. Together, our findings suggest that a setting-driven, tightly orchestrated cytotoxic-regulatory environment limits inflammation and favors *Mtb* control. Our results advocate for the use of TB endemic specimens to identify protective immune traits that allow tailoring TB treatment and vaccine development to specific settings and human populations.

## RESULTS

### *Mtb* replication is controlled significantly better by ex vivo granulomatous responses from healthy African compared to European PBMC donors

To investigate the impact of geographical origin on ex vivo infection outcomes, we applied a three-dimensional in vitro granuloma model of *Mtb* infection^21,22^ to PBMC specimens of healthy individuals recruited in Tanzania (n = 12) and at Swiss blood banks (n = 12) (**Fig. 1A**, Fig. S1). GSAv3-MD microarray typing confirmed predominantly European (mean proportion 0.93 ± 0.02 SD) and negligible African ancestry (mean 0.01 ± 0.01 SD) among the anonymous blood bank donors (**Fig. 1B**). We previously reported high genetic diversity among African ancestry subgroups in coastal Tanzania, alongside a minor contribution of non-African ancestry (mean 5%).^23^ We included two Tanzanian specimens in this ancestry typing using a different microarray to confirm minor European ancestry among the Tanzanian study participants (Fig. 1B). Self-reported tribe origin of African participants is provided in Supplementary Table 1. We will henceforth refer to European (EUR) for Swiss blood bank samples and African (AFR) for Tanzanian blood specimens. *Mtb* replication within ex vivo granulomas was variable (mean CFU/well 1.99×10^5^ ± 1.03×10^5^ SD; **Fig. 1C**), but consistent within individual donors across multiple experiments over time (**Fig. 1D**). Remarkably, PBMCs collected in AFR controlled *Mtb* growth significantly better than those collected in EUR did (**Fig. 1E**, *p* = 0.009).

**Figure 1.**
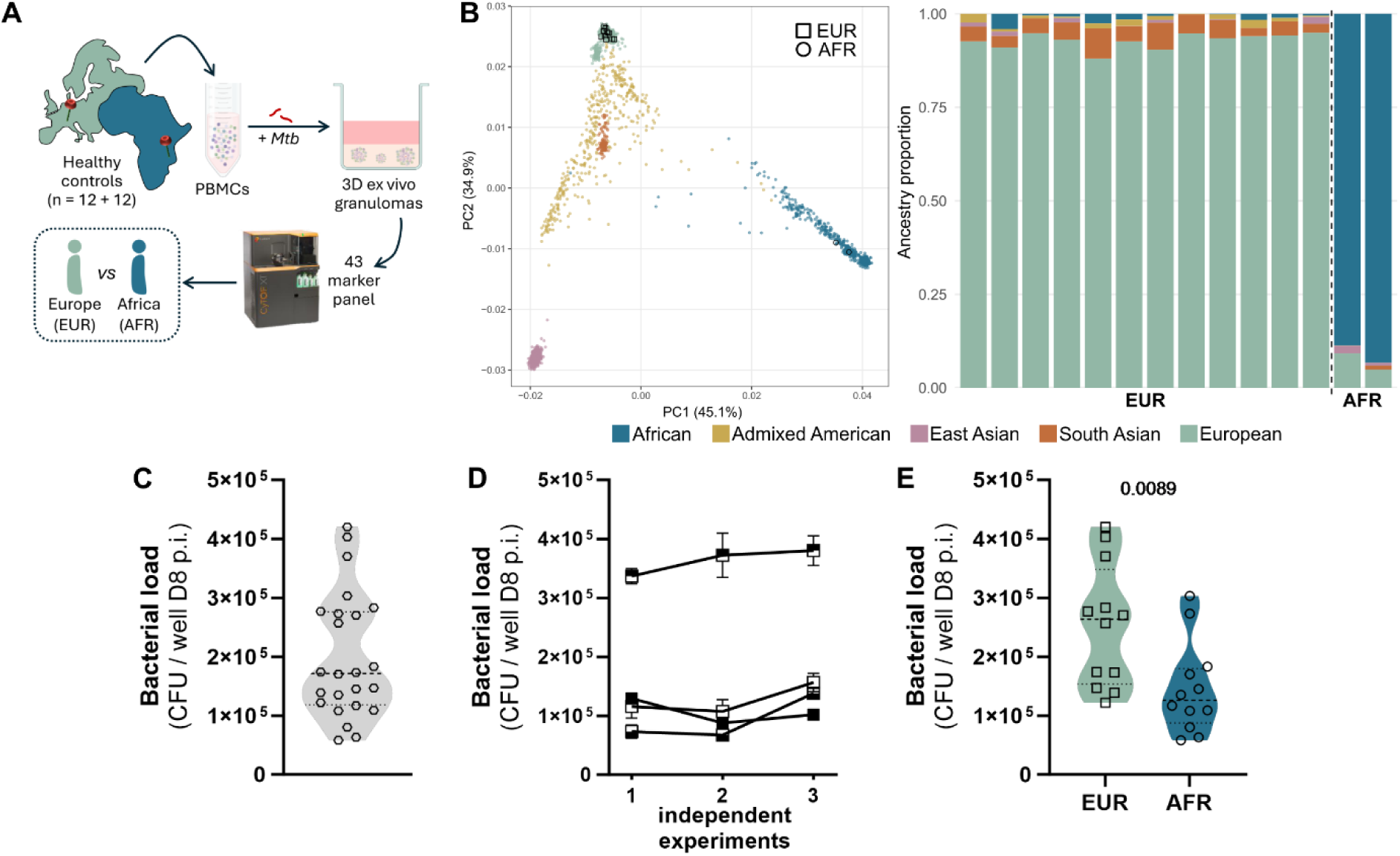
*Mtb* replication within human ex vivo granulomas is controlled significantly better by African PBMCs. (**A**) Schematic overview of the study design. (**B**) Ancestry typing (GSAv3-MD microarray) of EUR blood bank PBMCs (n=12) and Tanzanian reference PBMCs (n=2). PCA showing the localization of PBMCs among the 1000 Genomes Phase 3 reference panel (left) and bar plot depicting the ancestry proportion of each specimen (right). (C-E) *Mtb* load retrieved from granulomas eight days post-infection (D8 p.i.), quantified in colony-forming units (CFU): (**C**) Across all samples, independent of origin (n=24). (**D**) Reproducibility of *Mtb* load in granulomas from four independent donors (indicated in different symbols) across three independent experiments, each conducted two weeks apart. Error bars indicate SEM of technical replicates within each experiment. (**E**) Data from C split by sample origin (Europe (EUR): n=12; Africa (AFR): n=12); inset *p* value from an unpaired t-test.

### Deep immune profiling of ex vivo granulomas by CyTOF characterizes 728 functional phenotypes that differ by geographical origin

We next exploited a high-dimensional cytometry by-time-of flight (CyTOF) approach to unravel the cellular subsets that may confer *Mtb* control ex vivo within the granularity of 3D granulomas. We analyzed the combinatorial expression of 41 markers (Table 2) at an early (day 1; D1) and late (day 6; D6) time point post-infection. Employing FlowSOM^24^ clustering based on the expression of 28 canonical markers, we defined 20 lymphoid and 6 myeloid clusters (**Fig. 2A**). The highest proportion of cells were T (mean D1: 82.5%, D6: 86.8%), followed by B (D1: 9.1%, D6: 9.5%), NK (D1: 7.5%, D6: 3.4%), and myeloid cells (D1: 0.9%, D6: 0.3%). Uniform manifold approximations and projections (UMAPs) of the clusters across both groups visualized the contraction of the myeloid fractions, and expansion of certain T cell subsets (e.g. subset 11) over time (**Fig. 2B**). EUR and AFR granulomatous responses differed in cellular composition at both time points (PERMANOVA, D1: R^2^ = 0.185, *p* = 1×10^-4^; D6: R^2^ = 0.142, *p* = 4×10^-4^), with several differentially abundant subsets (**Fig. 2C**). Weak to moderate, positive and inverse correlations of subset frequencies with *Mtb* load were observed, but none of them reached statistical significance (**Fig. 2D**). Thus, we evaluated whether the cell subsets differed in function based on the expression of 28 functional markers rendering 728 potential cell subset-functional marker combinations (functional phenotypes) on day one (**Fig. 2E**) and day six (Fig. S2A). Functional phenotypes common to all samples (D1: n = 722, D6: n = 713) did not segregate based on bacterial load in a principal component analysis (PCA, Fig. S2B-C). However, PC1 on D1 (**Fig. 2F**) and PC2 on D6 (Fig. S2D) readily separated specimens based on geographical origin. Given that at least 230 functional phenotypes contributed significantly to the PCAs (Fig. S2E), we determined the most differential variables by calculating the magnitude of the group difference (Hedge’s g). At a significance threshold of *p*<0.001, AFR granulomatous responses on D6 were particularly enriched in granzyme B expressing (γδ) T cell 7 cells (Fig. S2F, S4A). On D1, we detected eleven differential functional phenotypes. AFR profiles were significantly enriched in activated and cytotoxic T cell subsets, particularly in perforin and granzyme B-expressing (γδ) T cell 7, CD127 (IL7-Ra)- and CD38-positive (CD8) T cell 4, CD127 (IL7-Ra) (CD4) T cell 12, as well as perforin expressing (CD8) T cell 3 cells. In contrast, EUR specimens harbored a higher frequency of pro- inflammatory myeloid cells, specifically TNF-α-, IL-6-, GM-CSF- and CD38-positive myeloid 2, as well as CD27-expressing (γδ) T cell 5 cells (**Fig. 2G**, Fig. S3B-C). Eight of the differential functional phenotypes showed significant inverse and positive correlations with bacterial load (**Fig. 2H**). Moreover, myeloid and lymphoid subsets displayed strong inverse correlations amongst themselves (Fig. S3D, Table 2).

**Figure 2.**
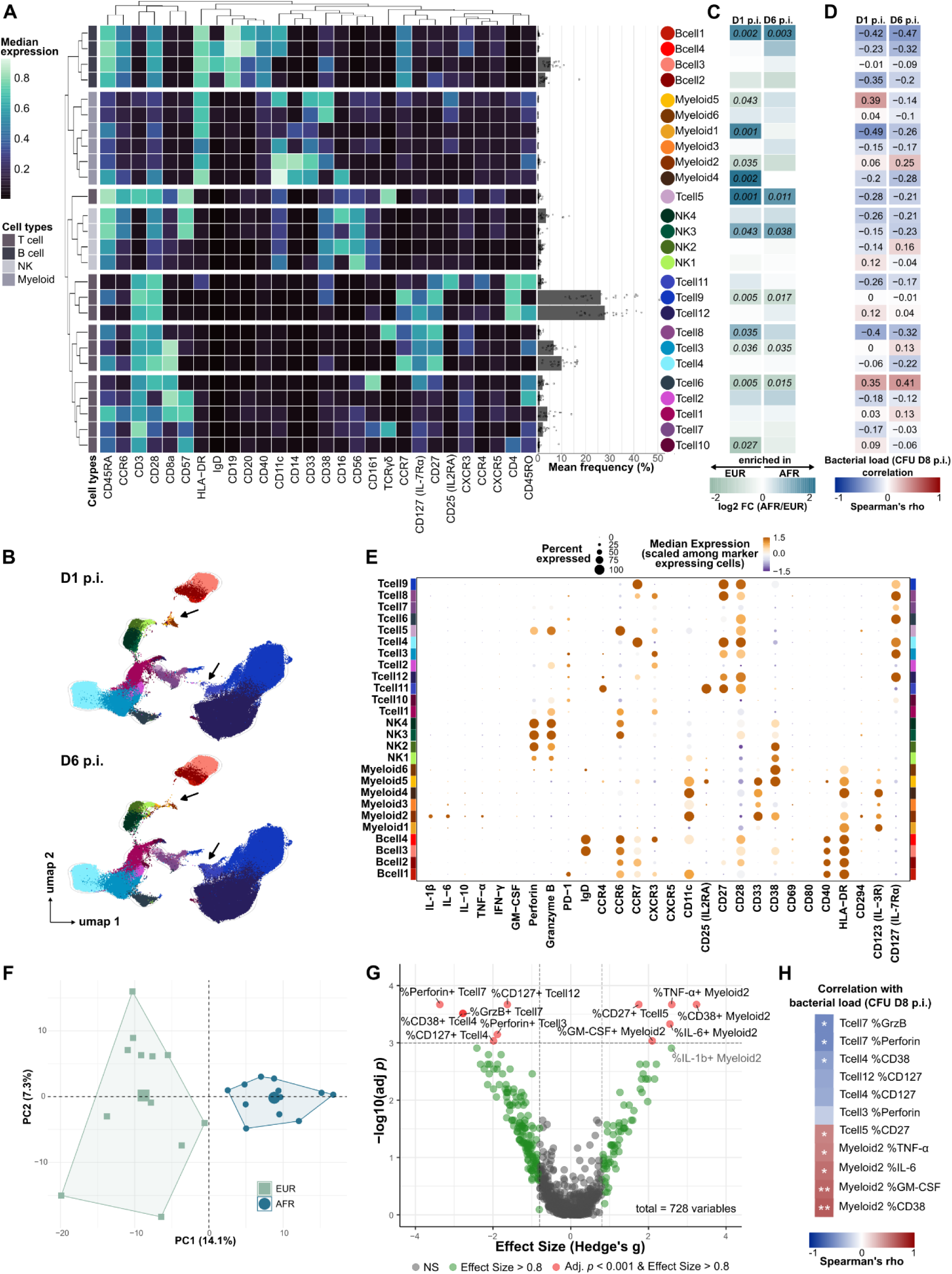
Deep immune profiling of ex vivo granulomas by CyTOF characterizes 728 functional phenotypes that differ by geographical origin. (**A**) Heatmap of median expression (D1 and D6 post-infection (p.i.) combined) of markers used for FlowSOM clustering. Barplot of mean frequency of each subset identified with FlowSOM; AFR (open circles, n=12), EUR (open squares, n=12) (**B**) UMAP visualization of FlowSOM subsets split by time point. (**C**) Differential abundance of subsets between EUR and AFR based on a negative binomial generalized linear model (GLM) using quasi-likelihood F-test. Color indicates log2 fold change (FC); inset *p* values are Benjamini-Hochberg corrected. (**D**) Spearman correlation of subset frequencies on D1 and D6 p.i. with bacterial load (CFU) retrieved on D8 p.i. Color and numbers indicate correlation coefficients. (**E**) Functional marker expression (D1 p.i.) by the different subsets. (**F**) PCA of data represented in Figure 2E, color-coded by country of origin. (**G**) Volcano plot (x: Hedge’s g effect size, y: false discovery rate (FDR)-adjusted *p*-values of pairwise Wilcoxon rank sum tests) of functional marker expression on D1 p.i.. (**H**) Spearman’s correlation analysis of D1 differential functional phenotypes with day eight bacterial load. Inset asterisks are FDR-corrected *p*-values. Figures E-G of the D6 data can be found as Supplementary Figures 3A, D, F.

### The EUR myeloid compartment is characterized by a strong inflammatory response to mycobacteria and LPS

Four of the five functional phenotypes enriched in EURs were myeloid 2 cells, a pro-inflammatory monocyte subset (Fig. S3C). The differential myeloid composition between EUR and AFR was visually apparent (**Fig. 3A**) and group accounted for 36% of the variance (PERMANOVA: R^2^ = 0.363, *p* = 1×10^-4^; Table 3) with no evidence of unequal dispersion. The myeloid 2 subset, a population significantly enriched in EURs (log2FC: -0.81, BH-adjusted *p* = 0.008), shifted in the UMAP with HLA-DR expression (**Fig. 3B**). While EUR myeloid 2 cells expressed higher levels of pro- inflammatory markers IL-1ϐ, IL-6, TNF-α, GM-CSF and CD38, HLA-DR expression was higher in AFR myeloid 2 cells (**Fig. 3C**, Fig. S3C). HLA-DR expression also characterized myeloid 1 cells, which AFRs were markedly enriched in (log2FC: 2.47, BH-adjusted *p* = 6.8×10^-6^). A pronounced pro-inflammatory response to infection, as displayed by the EUR myeloid subset, is reminiscent of innate immune training. We validated a stronger inflammatory response in EUR compared to AFR monocytes upon PBMC stimulation with lipopolysaccharide (LPS), *Mtb* soluble cell wall proteins (SCWP) and *Mtb* whole cell lysate (WCL) by flow cytometry. EUR monocytes produced significantly more IL-6 in response to all three stimuli and TNF-α in response to LPS (**Fig. 3D**, Fig. S4A). Corroborating our CyTOF data in an independent assay, the IL-6 response to mycobacterial stimulation in EUR monocytes also showed positive association with bacterial load in granulomatous responses, indicating that an excessive inflammatory response may cause a “cytokine storm” that favors bacterial growth (**Fig. 3E**). In contrast, the lower responsiveness of AFR monocytes may reflect innate immune tolerance^5,6^ supporting bacterial control. AFR monocytes have likely been trained through BCG vaccination (routinely administered in Tanzania at birth since 1975^25^, with coverage approaching 100%^26^), and setting-driven, repeated environmental exposure to mycobacteria and malaria parasites – all of which are known trained immunity inducers.^4,7,27^

**Figure 3.**
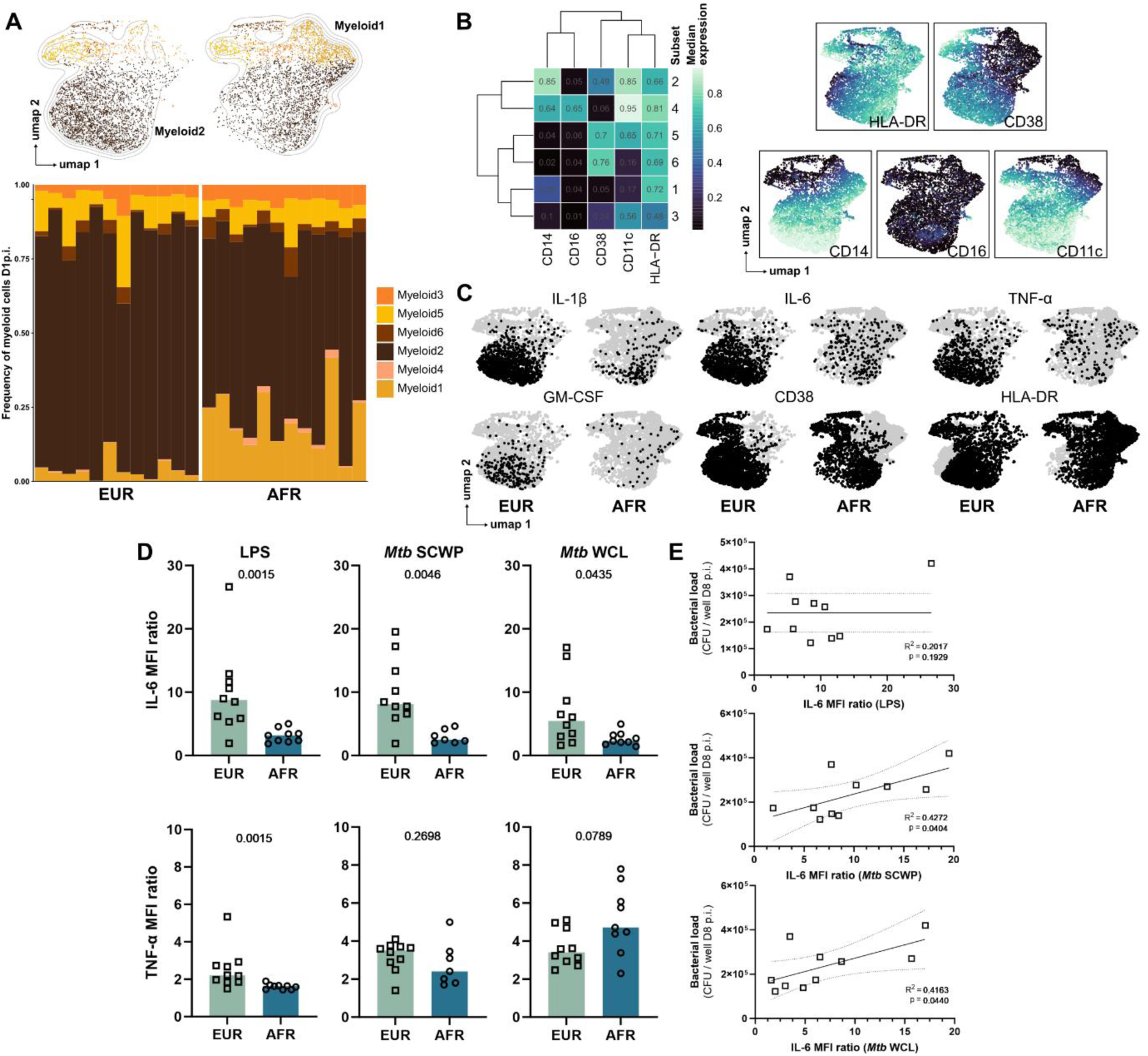
The EUR myeloid compartment is characterized by a strong inflammatory response to mycobacteria and LPS. (**A**) Composition of the myeloid compartment across origin visualized in a UMAP (*upper*) and as stacked bars (*lower*). (**B**) Heatmap and UMAP visualizations of the median expression of the clustering markers (both groups combined). (**C**) UMAP overlay of functional marker positive cells (black dots) and all cells (grey dots) split by origin. (**D**) Flow cytometry assessment (gating strategy provided as Fig. S4) of cytokine responses of monocytes to LPS, *Mtb* SCWP and *Mtb* WCL. Each dot represents an independent specimen (collected in Europe (EUR): n=10, or Africa (AFR): n=9; unless limited by sample availability that prioritized LPS read-out). Median fluorescence intensity (MFI) of IL-6 (*upper*) and TNF-α (*lower*) producing monocytes (ratio to mock control; inset *p* values indicate Mann-Whitney tests). (**E**) Linear fit of IL-6 MFI ratio in EUR monocytes and bacterial load in granulomas eight days post-infection (D8 p.i.). Statistical outlier detection was negative.

### Activated AFR T cells display higher polyfunctionality and cytotoxicity

Environmental exposure to mycobacteria may also shape adaptive immunity. Particularly, stronger T helper 1 (Th1) immunity could support the increased bacterial control observed in AFR granulomatous responses. Both EUR and AFR specimens displayed negligible canonical Th1 responses to stimulation with *Mtb*-specific peptides (ESAT-6/CFP-10/TB7.7) (**Fig. 4A**, Fig. S4B). In contrast, Th1 memory responses to stimuli containing cross-reactive antigens shared between *Mtb*, *M. bovis* BCG and environmental mycobacteria (SCWP and WCL) were significantly higher in AFR specimens (**Fig. 4B-C**). However, the increased frequency of mycobacteria-specific CD4 T cells did not correlate with a reduction of the bacterial load within respective ex vivo granulomas (**Fig. 4D**). This finding is consistent with our previous report^20^, in which AFR patients with TB did not control ex vivo *Mtb* infection better than AFR healthy controls (Fig. S5). Our CyTOF data revealed an enrichment in CD4, CD8 and γδ T functional phenotypes in AFR granulomatous responses characterized by activation and cytotoxicity and associated with reduced bacterial load (Fig. 2G-H). To further explore the differences in T cell functionality in EUR vs AFR granulomatous responses, we assessed activation and polyfunctionality (Special Polyfunctionality Index^28,29^, PI) of CD4, CD8 and γδ T cells on days 1 and 6 post-infection based on eight effector molecules (IL-1ϐ, IL-6, IL-10, TNF-α, IFN-γ, GM-CSF, perforin and granzyme B) across four different T cell activation definitions (**Fig. 4E**). Overall, AFR and EUR specimens showed comparable levels of T cell activation, except for CD8 and γδ T cells, which were more activated in AFRs in the most lenient definition. AFR CD4, CD8 and γδ T cells were more polyfunctional (cells with ≥ 2 functions) and showed higher PI in at least one subset-activation definition combination. Three subset x activation definition strata consistently showed higher PI medians and polyfunctionality distributions shifted towards higher degrees in AFRs across both time points (**Fig. 4E-F**). Multivariant marker intersections revealed that these strata were markedly enriched in granzyme B and perforin double-positive cells in AFR granulomatous responses (**Fig. 4G**). The frequency of granzyme B and perforin double-positive CD8 T cells (late and stringent activation definitions, D1 and D6) showed inverse correlation with bacterial load on day eight post-infection (**Fig. 4H**). This protective effect on bacterial control appeared to be group independent (partial Spearman correlation). Interestingly, the fourth most frequent marker combination in day 6 CD8 T cells (late and stringent) was co-expression of cytotoxic granzyme B, perforin and regulatory IL-10, which also showed inverse correlation with bacterial load (**Fig. 4I**).

**Figure 4.**
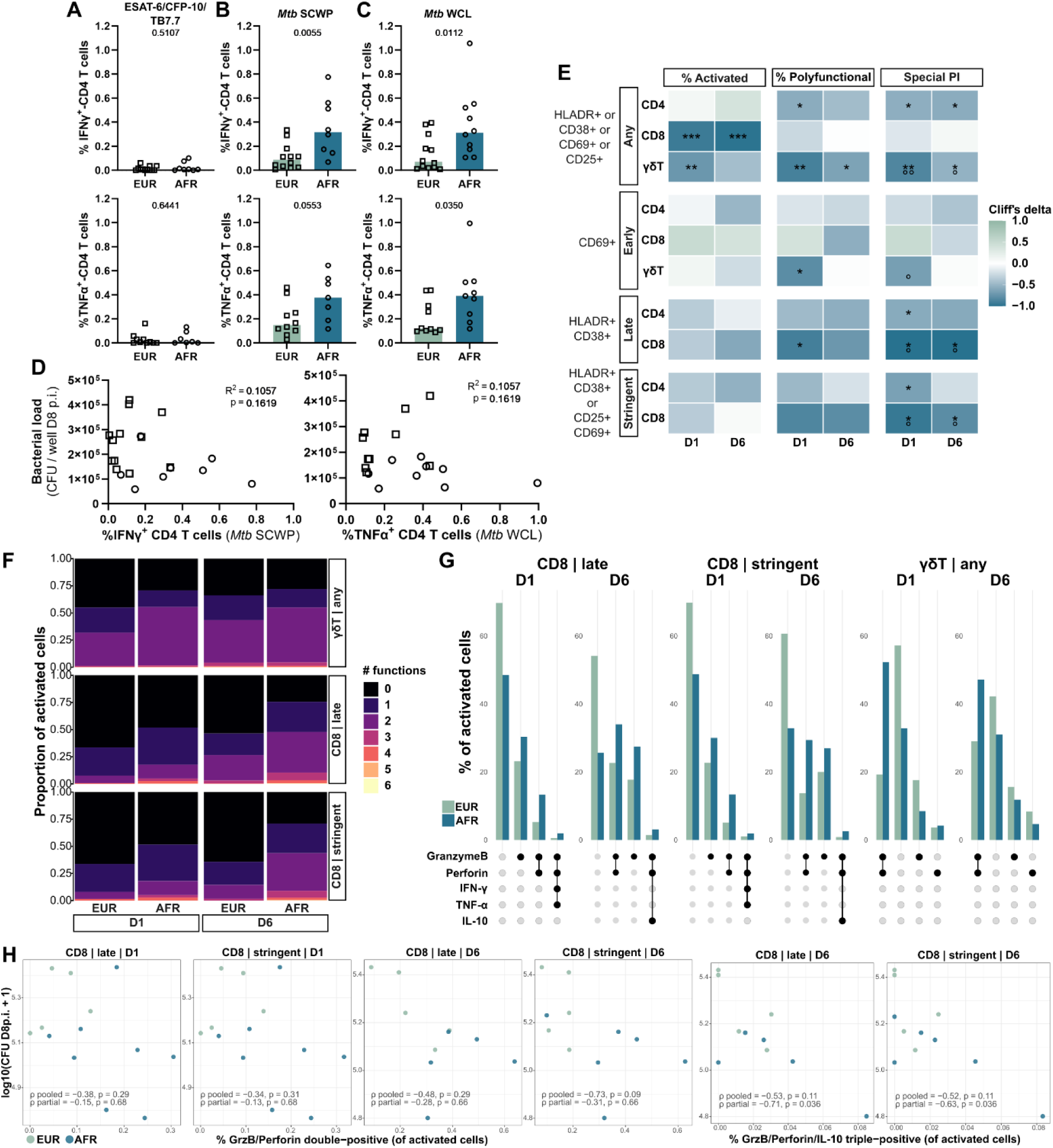
Activated AFR T cells display higher polyfunctionality and cytotoxicity. (**A-C**) Flow cytometry assessment of the frequency of IFN-γ- (*upper*) and TNF-α- (*lower*) positive CD4 T cells in response to (**A**) ESAT-6/CFP-10/TB7.7, (**B**) *Mtb* SCWP and (**C**) WCL. All percentages are background subtracted. Each dot represents an independent specimen (collected in Europe (EUR): n=12, or Africa (AFR): n=8). Inset *p* values indicate Mann-Whitney tests. (**D**) Linear fit of granuloma bacterial load on day 8 post-infection (D8 p.i.) with IFN-γ response to *Mtb* SCWP (*upper*) and TNF-α (*lower*), i.e. the most differential stimulation (of A-C) for each cytokine. (**E**) Assessment of T cell activation and polyfunctionality across four different activation definitions. Percent activated, percent polyfunctional (≥2 functions), polyfunctionality index (PI = mean[(#functions / 8)^q], q=1). Color shading indicates the effect size (Cliff’s delta; >0 higher in AFR); inset asterisks depict Wilcoxon *p* values FDR-adjusted per activation definition (Benjamini-Hochberg), inset degree symbols are of a L1 permutation test on the polyfunctionality distribution. (**F**) Visualization of polyfunctionality in strata with significantly different PI median (*) and distribution (°) between EURs and AFRs. (**G**) Upset plot visualizing the top 4 multivariant marker intersections of strata from F. (**H-I**) Spearman correlation (standard and partial accounting for group) of frequency of granzyme B, perforin double- (H) and granzyme B, perforin, IL-10 (I) triple-positive cells with log10(CFU D8 p.i.); inset *p* values are FDR-corrected (Benjamini-Hochberg).

Taken together, our deep immune profiling of ex vivo granulomatous responses to *Mtb* infection revealed high divergence across two geographical settings. The enrichment in highly cytotoxic and partially regulatory CD8 T cells, co-prevalent with an anti-inflammatory monocyte compartment in AFR responses, suggests a cytotoxic-regulatory environment limiting inflammation and favoring control of *Mtb* growth. We propose that this monocyte/T cell signature constitutes a promising proxy of protective immunity for vaccine evaluation that warrants further evaluation across independent cohorts.

## DISCUSSION

The extent and implications of human immune diversity have become increasingly apparent and recognized. However, we have yet to fully incorporate this diversity in the design of human immunology research and intervention development, particularly in the field of infectious diseases. We report highly divergent immune responses to mycobacterial infection across two different geographical settings that had functional consequences on the ex vivo infection control of *Mtb*. Significantly higher monocyte counts and plasma IL-6 levels have been associated with slower sputum conversion in Eurasian but not African TB patients.^30^ Our data further corroborate previous findings of increased viral^13,15^ and bacterial^11^ pathogen control by African compared to European specimens, although the opposite^12^ has also been observed. Likewise, a stronger inflammatory response has previously been linked to European^12,15^ and African^11,13^ ancestry, associated with increased^11–13^ or decreased^15^ pathogen control. While distinct pathogens and experimental inconsistencies may underlie part of these discrepancies, they also highlight how nuanced protective signatures may be. Moreover, immune responses to pathogens already differ at a smaller scale, within continents and even countries. In a cohort of admixed Peruvians, TB progression risk increased with native Peruvian genetic ancestry.^31^ Ex vivo, we have previously reported differential *Mtb*-specific CD4 T cell responses in Tanzanian compared to South African patients with active TB.^32^ Expression quantitative trait loci (eQTLs) have been identified as major contributors to ancestry-associated differences in immune responses to pathogens.^11,12,14^ However, inherited genetic variation and environmental influences are not readily separable, as both can shape epigenetic and transcriptional states that determine subsequent immune responsiveness. Recent work further demonstrates that epigenetic variation can contribute to inter-individual differences in infection responses.^33^ Thus, ancestry-associated immune phenotypes likely reflect a complex interplay between genetic, epigenetic, and environmental determinants.

Our approach has limitations. The sample size leaves the study underpowered to identify protective traits within each geographical setting. Moreover, the anonymity of the EUR specimens impeded controlling for confounding factors such as age and BCG vaccination status. Across our entire cohort, *Mtb* susceptibility may further be confounded by CMV co-infection, which largely accounted for ancestry-related differences in PBMC composition and responses to SARS-CoV-2.^14^ However, we previously showed that – similarly to EUR settings – CMV infection is predominantly acquired before adulthood in this AFR setting.^34^ Finally, ex vivo granulomas cannot fully reproduce in vivo pathogenesis of *Mtb* infection. They notably lack stromal and vascular components and do not reflect the chronic aspect of TB pathogenesis. Nevertheless, the comparison of ex vivo responses derived from human specimens of diverse genetic, immunological and environmental backgrounds captured differential immune responses directly affecting bacterial control.

These findings may inform the emerging concept of immune endotypes to guide host-directed therapies in TB^35^, as genetic background may represent an additional layer underlying distinct immune endotypes that could influence the response to immunomodulatory interventions. Moreover, in contrast to an emerging dogma suggesting that trained immunity-inducing vaccine adjuvants could increase their efficacy^36,37^, our data suggest that immune training may occur to the level of immune tolerance in AFR. In that context, inducing innate immune training is unlikely to improve TB vaccine efficacy in that particular setting. In sum, our data strongly support the need for holistic approaches that consider genetic, immunological and environmental differences to identify protective mechanisms in TB.

## MATERIALS AND METHODS

### PBMC preparation

European (EUR) peripheral blood mononuclear cell (PBMC) specimens were obtained under informed consent from healthy donors at the blood banks in Bern and Basel, Switzerland. African (AFR) PBMCs were obtained from healthy Tanzanians recruited within a diagnostic study conducted in Tanzania (TBChild project, IP.2009.32040.007), under a study protocol approved by the institutional review board of the Ifakara Health Institute (IHI-IRB), the Medical Research Coordinating Committee of the National Institute of Medical Research (NIMR) in Tanzania and the Ethikkommission Nordwest-und Zentralschweiz (EKNZ) in Switzerland. Ficoll-Paque^TM^ density-gradient centrifugation of fresh blood (AFR) and buffy coats (EUR) specimens was used to isolate PBMCs that were subsequently washed twice and resuspended in RPMI-1640 medium with L-glutamine (RPMI), supplemented with 10% DMSO and 40% heat-inactivated fetal calf serum (FCS), before aliquoting and cryopreservation in liquid nitrogen. According to need, cryopreserved PBMCs were thawed and washed twice in RPMI + 10% FCS + 12.5 U/ml benzonase, before overnight resting at 37°C, 5% CO2. PBMC viability above 95% was confirmed by trypan blue exclusion.

### Ancestry typing

Genomic DNA was isolated (Qiagen, #69504) from PBMCs and concentrated (Zymo, #D4010) as necessary prior genotyping at iGE3, Geneva, using the Illumina GSA v3-MD array (GRCh37/hg19) and Illumina GenomeStudio (v2, Illumina, Inc.) for genotype calling. Quality control (SNP call rate >95%, sample missingness filter, sex check, heterozygosity check, Hardy-Weinberg equilibrium) was performed in PLINK v1.9^38^, leading to a final dataset of 689,632 SNPs across all samples. To infer continental ancestry, samples were merged with 2,318 individuals from the 1000 Genomes Project Phase 3 reference panel^39^ genotyped on the Illumina Omni2.5 array, yielding 125,505 shared autosomal SNPs. Following linkage disequilibrium (LD) pruning using a sliding window approach (window size 50 SNPs, step 10, r^2^ threshold 0.1), principal component analysis (PCA) was performed on the 46,092 retained SNPs to assess population structure relative to the reference superpopulations using PLINK v1.9. Unsupervised ancestry estimation was done using ADMIXTURE^40^ (v1.3.0), with the number of ancestral clusters (K) ranging from 2 to 6. The optimal K was selected based on 10-fold cross-validation error, which plateaued at K=5 (CV error = 0.563), corresponding to the five continental reference superpopulations. Bioinformatics pipeline development was assisted by Claude (Anthropic).

### Mycobacterial strain preparation

*Mtb* H37Rv grown to mid-exponential phase (OD_600_ ≈ 0.7) in Middlebrook 7H9 broth supplemented with 10% ADC (5% bovine albumin fraction V, 2% dextrose and 0.003% catalase), 0.5% glycerol and 0.05% Tween-80 was used to obtain a suspension of single-cell bacilli as follows. Bacteria were pelleted at 3000×g (5 min), washed in PBS + 0.1% Tween-80 and re-suspended in RPMI + 10% FCS. The bacteria were subsequently dispersed for two minutes in a water-bath sonicator (XUB5, Grant Instruments), followed by a centrifugation at 260×g (5 min) to pellet remaining clumps. The upper part of the supernatant was supplemented with 5% glycerol final before aliquoting and storage at -80°C and its concentration determined by colony forming units (CFU) on Middlebrook 7H11 supplemented with 10% ADC + 0.05% oleic acid (OADC) upon serial dilution in PBS-Tween-80 (0.1%).

### 3D in vitro granuloma model

Human PBMCs were infected and embedded in an extracellular matrix to allow spontaneous formation of granulomatous structures precisely as previously reported.^20^ In brief, PBMCs were infected with H37Rv (MOI 1:400), embedded and topped up with RPMI+20% HuS for eight days. Granuloma formation was monitored by bright-field microscopy and supernatants exchanged after 4 days. Following overnight brefeldin A treatment, host cells were retrieved on days 1 and 6 post-infection for subsequent CyTOF staining. Retrieved host cells were lysed for CFU enumeration on day 8 post-infection.

### CyTOF staining, acquisition and analysis

The samples used in the study (EUR n = 12, AFR n = 12) were stained and acquired jointly with TB patient samples (n = 11), as previously reported.^20^ Data pre-processing (bead normalization^41^, compensation^42^, de-barcoding^43^ and manual gating (FlowJo 10.6.1), batch correction^44^ and arcsinh-transformation) was performed in parallel as described.^20^ Subsequently, the dataset was split to assess the effect of disease status (AFR healthy vs patients, previously reported^20^) and geographical origin (AFR vs EUR healthy, present study), respectively. All downstream analyses were performed in R (version 4.4.1). Data presented here are based on 4,570,301 cells from 24 independent specimens across two time points. In brief, cells were clustered in FlowSOM^24^ using a hierarchical approach that first split the data into major cell types (T, B, NK and myeloid cells) based on canonical lineage markers (CD3, CD4, CD8a, CD19, CD20, HLADR, CD56NCAM, CD16, CD14, CD11c, CD33, CD40), a 10x10 grid and 15 metaclusters. Secondly, clustering within each cell type was performed: for T cells markers CD3, CD4, CD8a, CD27, CD28, CD45RA, CD45RO, CD56, CD161, TCRgd, CCR4, CCR6, CXCR3, CXCR5, a 10x10 grid and 12 meta clusters; for B cells markers CD11c, CD19, CD20, CD27, a 8x8 grid and 4 meta clusters; for NK cells markers CD56, CD57, CD16, a 8x8 grid and 4 meta clusters; and for myeloid cells markers CD11c, CD14, CD16, CD38, HLA-DR, a 8x8 grid and 6 meta clusters. Data were visualized using “umap”^45^. Marker positivity was estimated per sample using silhouette-scanning in “MetaCyto”^46^ and overall median cutoffs were used to guide manual corrections. Marker expression was visualized in bubble plots using “Seurat”.^47^ PCA analyses were performed in “factoextra”.^48^ and volcano plots in “EnhancedVolcano”^49^ using previously published functions.^50^ Differences in overall cell composition between groups was assessed by PERMANOVA (vegan::adonis2) using Euclidean distances on centered log-ratio (CLR) transformed subset frequencies (pseudocount replacement for zeros and 9,999 permutations) to account for their compositional nature. Homogeneity of multivariate dispersion was confirmed using betadisper. To assess differential abundance of individual subsets between groups quasi-likelihood negative binomial generalized linear models implemented in “edgeR” were used following the diffcyt^51^ workflow. Raw cell counts were modeled for each subset, with the total number of cells per sample used as the library size. Library sizes were normalized using trimmed mean of M-values (TMM) method, and empirical Bayes quasi-likelihood F-tests were used to assess differential abundance between groups. Polyfunctionality was defined as cells expressing ≥2 effector molecules (IL-1ϐ, IL-6, IL-10, TNF-α, IFN-γ, GM-CSF, perforin and granzyme B). The previously defined special polyfunctionality index (PI)^28,29^ was calculated across all eight effector molecules, assuming equal contributions and setting q = 1, such that PI represents the mean fraction of functions engaged per activated cell. PI was calculated per donor for each T cell subset (CD4, CD8 and γδ), timepoint and activation definition.

### Host cell stimulation

Rested PBMCs (2.5×10^5^ cells per condition in 200 μl final volume of RPMI + 10% FCS) were incubated with LPS (1 μg/ml), *Mtb* whole cell lysate (10 μg/ml; NR-14822, BEI Resources), *Mtb* peptide pool covering ESAT-6/CFP-10/TB7.7 (5 μg/ml), *Mtb* soluble cell wall proteins (10 μg/ml; NR-14840, BEI Resources), or medium (unstimulated control). After 8 hours, half of the supernatant was replaced with RPMI + 10% FCS and brefeldin A (5 μg/ml). PBMC stimulation was pursued for another 15 h before staining for flow cytometry. In brief, PBMCs were stained extracellularly with CD14 (QA18A22)-APC/Cy7 antibody (Biolegend) for 15 min at room temperature, fixed and permeabilized (Biolegend, following manufacturer’s protocol). Intracellular staining with anti-CD3 (UCHT1)-PB, anti-TNF-α (Mab11)-BV510, anti-CD4 (RPA-T4)-PE, anti-IFN-γ (4S.B3)-PerCP and anti-IL6 (MQ2-13A5)-PE/Cy7 antibodies (Biolegend) was performed during 30 min at room temperature before washing and acquisition on a MACSQuantX instrument (Miltenyi Biotec).^23^

## ACKNOWLEDGMENTS AND FUNDING SOURCES

We are grateful to the TBCHILD project (IP.2009.32040.007) study participants, funded by the European and Developing Countries Clinical Trials Partnership (EDCTP) and donors at the blood banks of Bern and Basel, Switzerland for consenting to donate their blood for research purposes. We acknowledge the Swiss TPH BSL3 staff for their technical support. This research was funded by the Swiss National Science Foundation (grant numbers 197838 and 177163). Computational analyses were performed at sciCORE (https://scicore.unibas.ch) scientific computing center at University of Basel.

