## Supplementary for "Deep CyTOF immune profiles of ex vivo granulomatous responses to *Mycobacterium tuberculosis* differ with European and African geographical origin"

### SUPPLEMENTARY MATERIALS

#### Supplementary figures

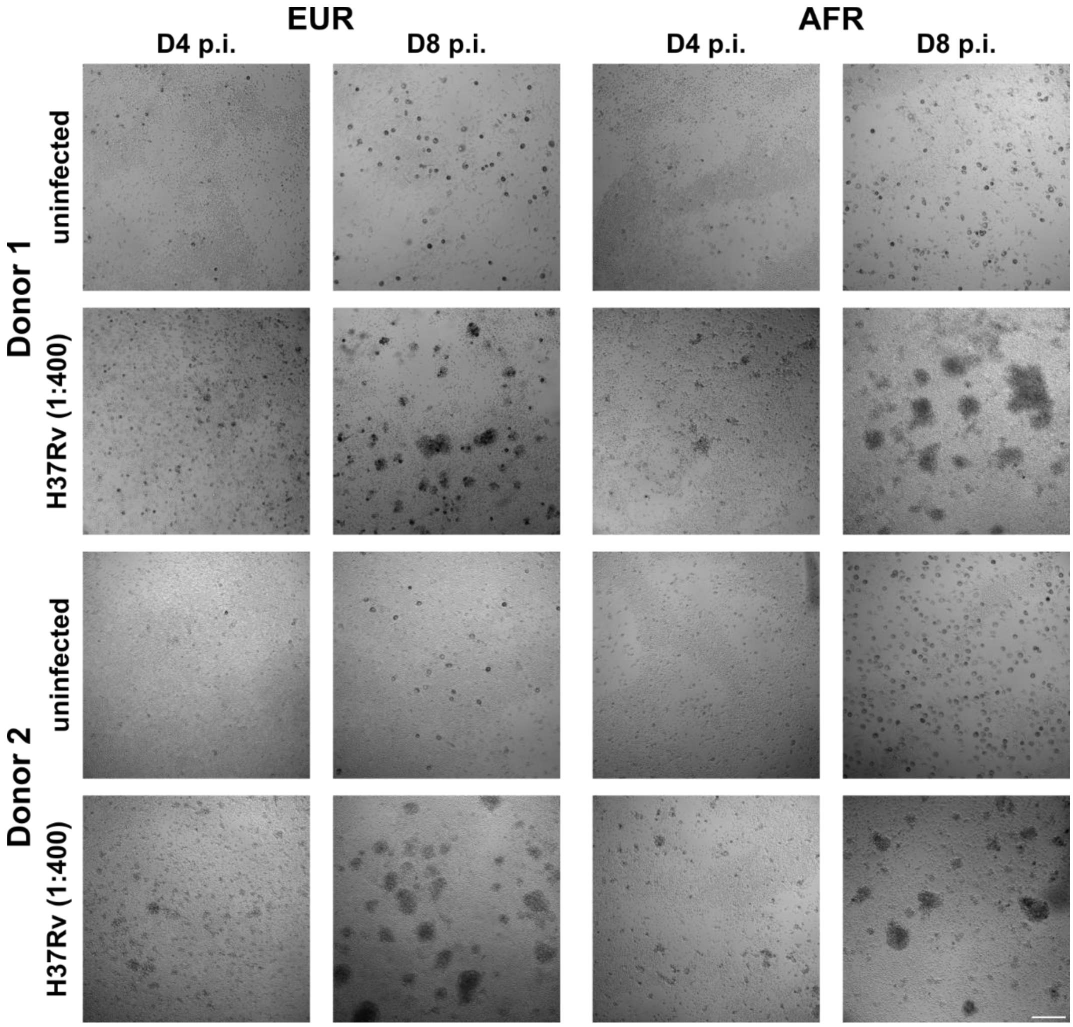


**Figure S1.** Pictures of granuloma formation by bright field microscopy four (*left*) and eight (*right*) days post-infection (p.i.) and matrix embedding of four representative PBMC specimens used in this study, two of each geographic origin. Scale bar is 200 µm.

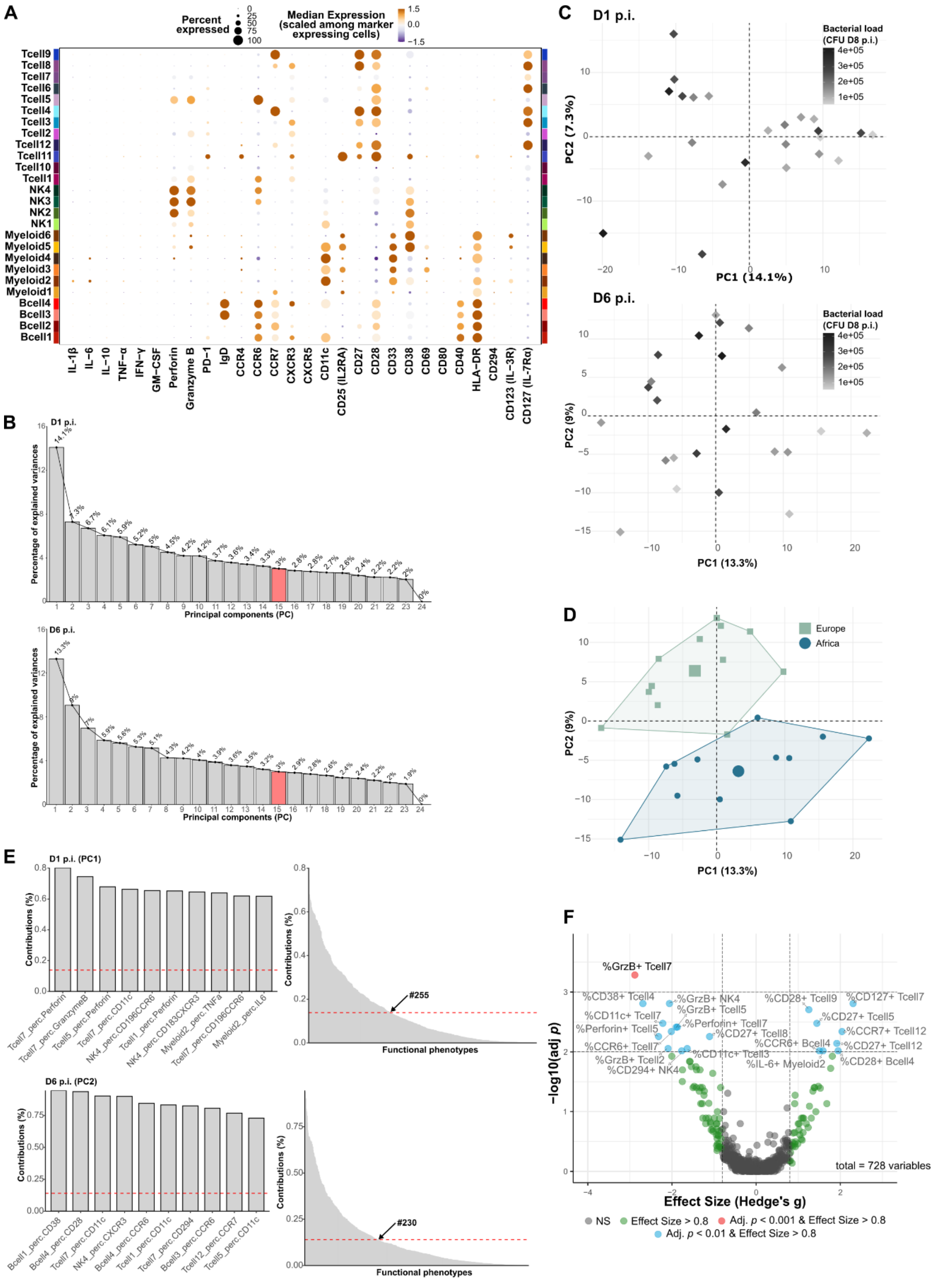


**Figure S2. Functional marker expression on FlowSOM subsets and PCA**. (**A**) Functional marker expression by the different cell subsets on day 6 post-infection (D6 p.i.). (**B**) Scree plots of day one (*upper*) and day six (*lower*) PCAs of functional phenotypes. Filled red bar indicates number of PCs needed to account for 80% of the data sets' variances. (**C**) D1 and D6 PCA color-coded by bacterial load (CFU) retrieved on D8 p.i. (**D**) D6 PCA color-coded by country of origin. (**E**) Contributions to PC1 on D1 (*upper*) and PC2 (*lower*) on D6 p.i.. Left plots highlight the top ten contributing functional phenotypes for the indicated PC and time point p.i.. The plots on the right show the contribution of all functional phenotypes to a given PC and time point. The arrow indicates the number of functional phenotypes showing contributions above the expected contribution (red dotted line). The expected contribution assumes equal contribution of each variable and is calculated as: 1/number of variables (i.e. functional phenotypes) ×100. (**F**) Volcano plot (x: Hedge's g effect size, y: false discovery rate (FDR)-adjusted *p*-values of pairwise Wilcoxon rank sum tests) of functional marker expression on D6 p.i..

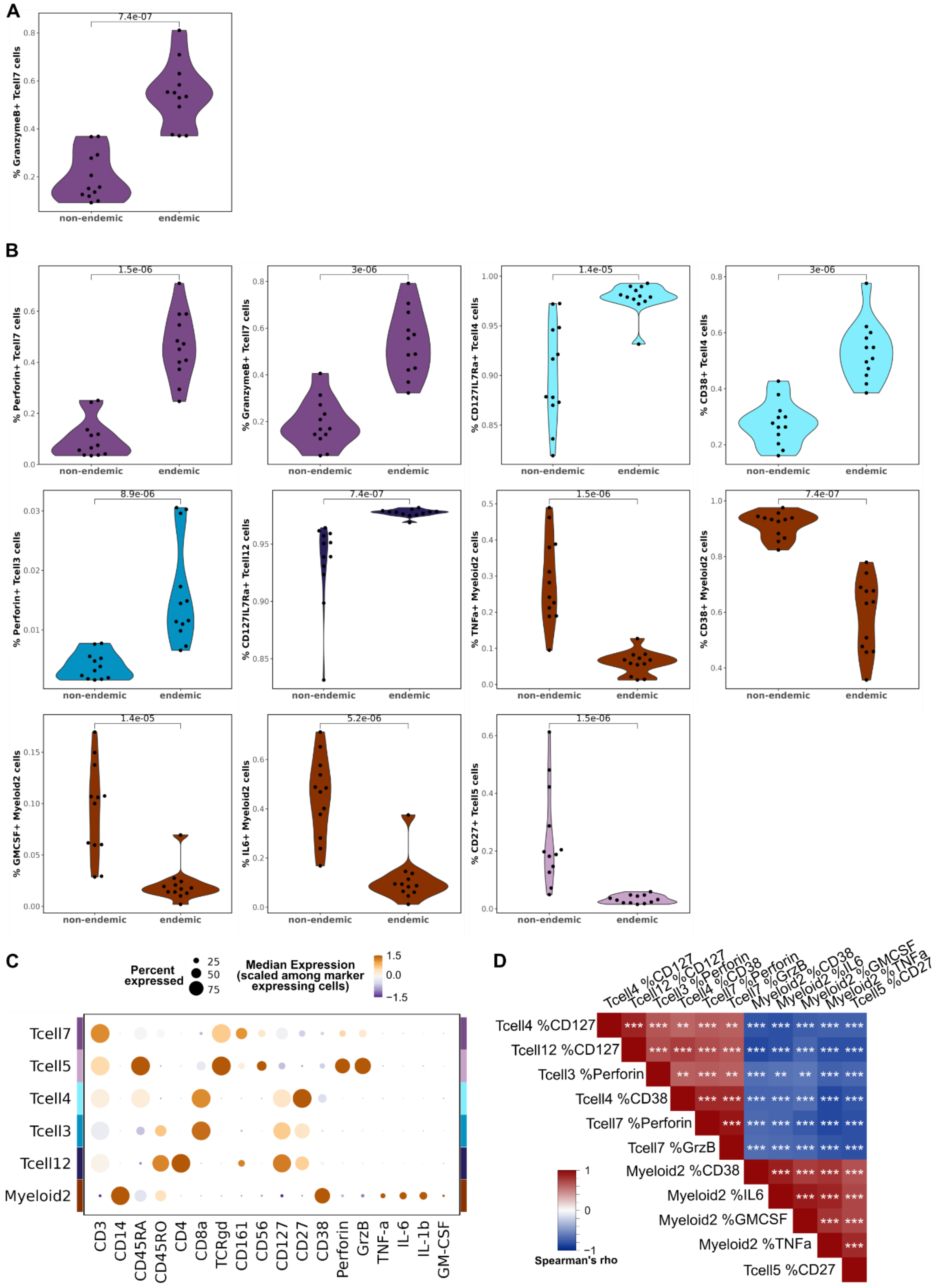


**Figure S3. Differential functional phenotypes in ex vivo granulomas.** Differential functional phenotypes on (**A**) day six and (**B**) day one post-infection (p.i.), as identified in Figures S3F and

2G. Inset *p* values are of Wilcoxon tests. (**C**) Percent expressed and median expression level of functional and canonical markers characterizing the subsets with differential functional phenotypes. (**D**) Spearman's correlation analysis of the day one differential functional phenotypes amongst themselves. Inset asterisks are FDR-corrected *p*-values.

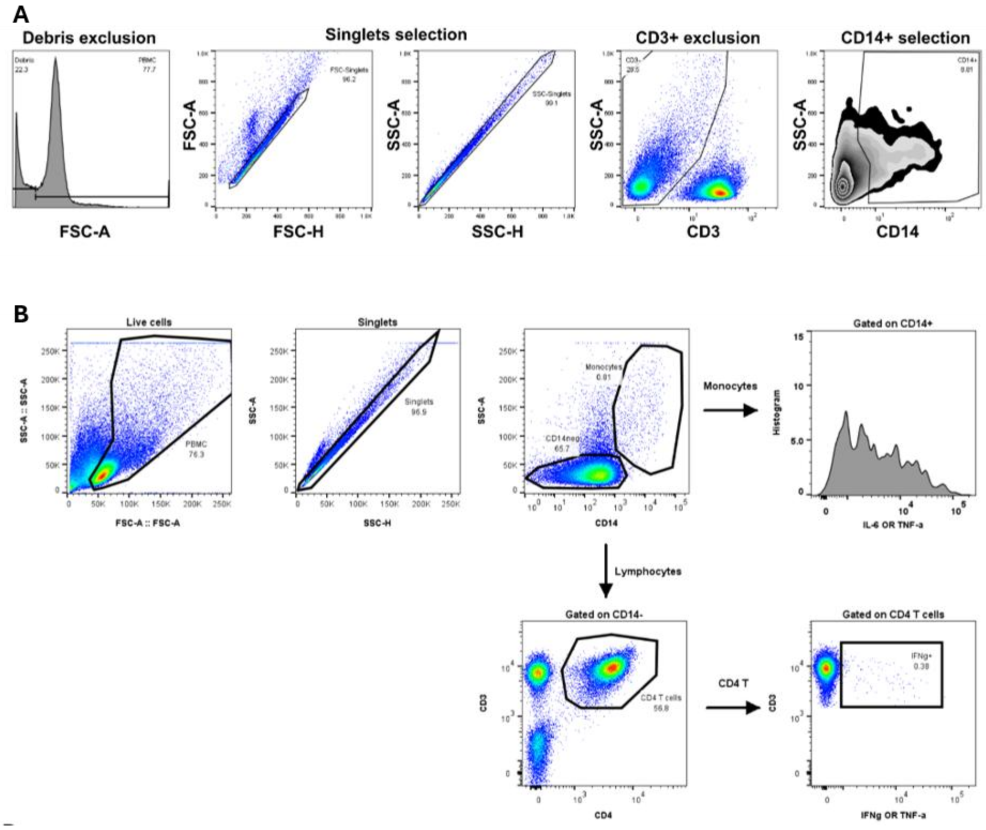


**Figure S4. Flow cytometry gating strategies.** (**A**) Data underlying figure 3D. (**B**) Data underlying figures 4A-C.

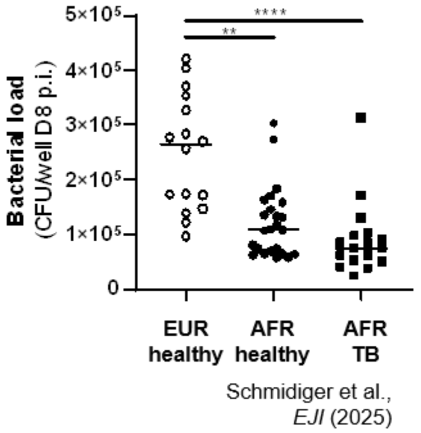


**Figure S5. Superior control of *Mtb* growth within ex vivo granulomas derived from AFR compared to EUR specimens is independent of TB disease status**

### Supplementary tables

#### Table 1: Specimen characteristics

| CHARACTERISTIC | | EUR (n=12) | AFR (n=12) |
|---|---|---|---|
| **Ancestry information** | Typed, n (%) | 12 (100) | 0 (0) |
| | Self-reported, n (%) | 0 (0) | 10 (80) |
| **Ancestry** | European, n (%) | 12 (100) | 0 (0) |
| | African, n (%) | 0 (0) | 12 (100) |
| | **Tribe affiliation** | N/A | |
| | *Wakwere* | | 1 (8.3) |
| | *Zaramo* | | 3 (16.7) |
| | *Gogo* | | 1 (8.3) |
| | *Mpare* | | 1 (8.3) |
| | *Doe* | | 1 (8.3) |
| | *Makonde* | | 1 (8.3) |
| | *Ngoni* | | 1 (8.3) |
| | *Haya* | | 1 (8.3) |
| | *NA* | | 2 (16.7) |
| **Sex**, n female (%) | | 5 (41.7) | 1 (8.3) |
| **Age (years)**, median (IQR) | | *NA* | 21 (4.75) |

*NA* data not available; N/A characteristic not applicable

#### Table 2: CyTOF marker panel

| Target (clone) | Label | Used for | |
|---|---|---|---|
| | | FlowSOM clustering | Functional assessment |
| **CCR4** (L291H4) | 152Sm | YES | YES |
| **CCR6** (G034E3) | 141Pr | YES | YES |
| **CCR7** (G043H7) | 167Er | YES | YES |
| **CD11c** (Bu15) | 147Sm | YES | YES |
| **CD123** (6H6) | 143Nd | | YES |
| **CD127** (A019D5) | 176Yb | YES | YES |
| **CD14** (63D3) | 168Er | YES | |
| **CD16** (3G8) | 148Nd | YES | |
| **CD161** (HP-3G10) | 151Eu | YES | |
| **CD19** (HIB19) | 144Nd | YES | |
| **CD20** (2H7) | 171Yb | YES | |
| **CD25** (BC96) | 153Eu | YES | YES |
| **CD27** (O323) | 154Sm | YES | YES |
| **CD28** (CD28.2) | 160Gd | YES | YES |
| **CD294** (BM16) | 166Er | | YES |
| **CD3** (UCHT1) | 170Er | YES | |
| **CD33** (WM53) | 169Tm | YES | YES |
| **CD38** (HB-7) | 161Dy | YES | YES |
| **CD4** (RPA-T4) | 145Nd | YES | |
| **CD40** ( 5C3) | 142Nd | YES | YES |
| **CD45RA** (HI100) | 150Nd | YES | |
| **CD45RO** (UCHL1) | 149Sm | YES | |
| **CD56** (NCAM16.2) | 163Dy | YES | |
| **CD57** (HNK-1) | 155Gd | YES | |
| **CD69** ( FN50) | 113Cd | | YES |
| **CD80** ( 2D10.4) | 162Dy | | YES |
| **CD8a** (RPA-T8) | 146Nd | YES | |
| **CXCR3** (G025H7) | 156Gd | YES | YES |
| **CXCR5** (J252D4) | 158Gd | YES | YES |
| **GM-CSF** (BVD221C11) | 159Tb | | YES |
| **Granzyme B** (GB11) | 198Pt | | YES |
| **HLA-DR** (LN3) | 173Yb | YES | YES |
| **IFN-γ** (B27) | 116Cd | | YES |
| **IgD** (IA6-2) | 174Yb | YES | YES |
| **IL-10** (JES3-9D7) | 165Ho | | YES |

| | | | |
|---|---|---|---|
| **IL-1β** (CRM56) | 209Bi | | YES |
| **IL-6** (MQ2-13A5) | 106Cd | | YES |
| **PD-1** ( EH12.2H7) | 175Lu | | YES |
| **Perforin** (B-D48) | 196Pt | | YES |
| **TCRγδ** (B1) | 164Dy | YES | |
| **TNF-α** (Mab11) | 114Cd | | YES |
| CD66b (G10F5) | 172Yb | *excluded from analysis* | |
| CD45 (HI30) | 089Y | *excluded after data clean-up* | |
| DNA1 | 191Ir | *used for data clean-up / manual gating* | |
| DNA2 | 193Ir | | |
| Live/dead | 103Rh | | |
| CD45 (HI30) *barcoding* | 110Cd | *used for multiplexing* | |
| | 194Pt | | |
| | 111Cd | | |
| | 195Pt | | |

### Table 2: Spearman correlations of volcano hits with CFU and amongst themselves

| Functional phenotype | Spearman rho | BH adjusted p |
|---|---|---|
| % CD38$^{+}$ Myeloid 2 | 0.644348 | 0.007457 |
| % GM-CSF$^{+}$ Myeloid 2 | 0.614783 | 0.007644 |
| % IL-6$^{+}$ Myeloid 2 | 0.567826 | 0.012489 |
| % Perforin$^{+}$ T cell 7 | -0.54696 | 0.012489 |
| % Granzyme B$^{+}$ T cell 7 | -0.54783 | 0.012489 |
| % TNF-α$^{+}$ Myeloid 2 | 0.508696 | 0.020415 |
| % CD27$^{+}$ T cell 5 | 0.484348 | 0.025874 |
| % CD38$^{+}$ T cell 4 | -0.45913 | 0.033021 |
| % CD127$^{+}$ T cell 12 | -0.40696 | 0.059186 |
| % CD127$^{+}$ T cell 4 | -0.38609 | 0.068637 |
| % Perforin$^{+}$ T cell 3 | -0.23217 | 0.274965 |

| Functional phenotype 1 | Functional phenotype 2 | Spearman rho | BH adjusted p |
|---|---|---|---|
| % Perforin$^{+}$ T cell 7 | % GranzymeB$^{+}$ T cell 7 | 0.938261 | 7.06E-10 |
| % IL-6$^{+}$ Myeloid 2 | % GMCSF$^{+}$ Myeloid 2 | 0.906957 | 2.77E-08 |
| % CD38$^{+}$ Myeloid 2 | % IL-6$^{+}$ Myeloid 2 | 0.893913 | 7.35E-08 |
| % CD127$^{+}$ T cell 12 | % CD127$^{+}$ T cell 4 | 0.873913 | 3.35E-07 |
| % IL-6$^{+}$ Myeloid 2 | % TNF-α$^{+}$ Myeloid 2 | 0.864348 | 5.72E-07 |
| % TNF-α$^{+}$ Myeloid 2 | % CD38$^{+}$ T cell 4 | -0.85565 | 9.04E-07 |
| % CD38$^{+}$ Myeloid 2 | % CD127$^{+}$ T cell 12 | -0.85217 | 9.22E-07 |
| % TNF-α$^{+}$ Myeloid 2 | % Perforin$^{+}$ T cell 7 | -0.84957 | 9.22E-07 |
| % CD38$^{+}$ T cell 4 | % Perforin$^{+}$ T cell 7 | 0.849565 | 9.22E-07 |
| % TNF-α$^{+}$ Myeloid 2 | % CD27$^{+}$ T cell 5 | 0.846957 | 9.89E-07 |
| % CD38$^{+}$ T cell 4 | % GranzymeB$^{+}$ T cell 7 | 0.845217 | 1.01E-06 |
| % TNF-α$^{+}$ Myeloid 2 | % GMCSF$^{+}$ Myeloid 2 | 0.82 | 3.97E-06 |
| % CD27$^{+}$ T cell 5 | % Perforin$^{+}$ T cell 7 | -0.82 | 3.97E-06 |
| % IL-6$^{+}$ Myeloid 2 | % CD127$^{+}$ T cell 4 | -0.8087 | 5.96E-06 |
| % CD127$^{+}$ T cell 12 | % CD27$^{+}$ T cell 5 | -0.8087 | 5.96E-06 |
| % CD27$^{+}$ T cell 5 | % GranzymeB$^{+}$ T cell 7 | -0.80957 | 5.96E-06 |
| % IL-6$^{+}$ Myeloid 2 | % CD127$^{+}$ T cell 12 | -0.80696 | 6.14E-06 |
| % CD38$^{+}$ Myeloid 2 | % TNF-α$^{+}$ Myeloid 2 | 0.804348 | 6.64E-06 |
| % TNF-α$^{+}$ Myeloid 2 | % CD127$^{+}$ T cell 12 | -0.80261 | 6.87E-06 |
| % CD38$^{+}$ Myeloid 2 | % CD127$^{+}$ T cell 4 | -0.78435 | 1.43E-05 |
| % CD38$^{+}$ T cell 4 | % CD27$^{+}$ T cell 5 | -0.78522 | 1.43E-05 |
| % TNF-α$^{+}$ Myeloid 2 | % GranzymeB$^{+}$ T cell 7 | -0.78609 | 1.43E-05 |

| | | | |
|---|---|---|---|
| % CD127$^{+}$ T cell 12 | % CD38$^{+}$ T cell 4 | 0.771304 | 2.45E-05 |
| % CD38$^{+}$ Myeloid 2 | % GMCSF$^{+}$ Myeloid 2 | 0.766087 | 2.93E-05 |
| % GMCSF$^{+}$ Myeloid 2 | % Perforin$^{+}$ T cell 7 | -0.76348 | 3.13E-05 |
| % GMCSF$^{+}$ Myeloid 2 | % CD127$^{+}$ T cell 4 | -0.74174 | 7.09E-05 |
| % TNF-α$^{+}$ Myeloid 2 | % Perforin$^{+}$ T cell 3 | -0.73739 | 8.02E-05 |
| % GMCSF$^{+}$ Myeloid 2 | %CD27$^{+}$ T cell 5 | 0.735652 | 8.24E-05 |
| % TNF-α$^{+}$ Myeloid 2 | %CD127$^{+}$ T cell 4 | -0.73217 | 9.03E-05 |
| % IL-6$^{+}$ Myeloid 2 | %CD27$^{+}$ T cell 5 | 0.729565 | 9.27E-05 |
| % CD127$^{+}$ T cell 4 | %CD27$^{+}$ T cell 5 | -0.72957 | 9.27E-05 |
| % GMCSF$^{+}$ Myeloid 2 | %CD127$^{+}$ T cell 12 | -0.71391 | 0.000141 |
| % CD38$^{+}$ Myeloid 2 | % CD38$^{+}$ T cell 4 | -0.71391 | 0.000141 |
| % IL-6$^{+}$ Myeloid 2 | % CD38$^{+}$ T cell 4 | -0.71652 | 0.000141 |
| % IL-6$^{+}$ Myeloid 2 | % Perforin$^{+}$ T cell 7 | -0.71478 | 0.000141 |
| % CD127$^{+}$ T cell 12 | % Perforin$^{+}$ T cell 3 | 0.708696 | 0.000162 |
| % CD127$^{+}$ T cell 12 | % Perforin$^{+}$ T cell 7 | 0.706087 | 0.000167 |
| % GMCSF$^{+}$ Myeloid 2 | % GranzymeB$^{+}$ T cell 7 | -0.70609 | 0.000167 |
| % Perforin$^{+}$ T cell 3 | % CD27$^{+}$ T cell 5 | -0.70435 | 0.000172 |
| % CD38$^{+}$ Myeloid 2 | % CD27$^{+}$ T cell 5 | 0.69913 | 0.000198 |
| % CD38$^{+}$ Myeloid 2 | % Perforin$^{+}$ T cell 7 | -0.69739 | 0.000204 |
| % IL-6$^{+}$ Myeloid 2 | % GranzymeB$^{+}$ T cell 7 | -0.68957 | 0.000253 |
| % CD38$^{+}$ Myeloid 2 | % Perforin$^{+}$ T cell 3 | -0.67565 | 0.000364 |
| % GMCSF$^{+}$ Myeloid 2 | % CD38$^{+}$ T cell 4 | -0.67565 | 0.000364 |
| % CD127$^{+}$ T cell 12 | % GranzymeB$^{+}$ T cell 7 | 0.673913 | 0.000374 |
| % CD38$^{+}$ Myeloid 2 | % GranzymeB$^{+}$ T cell 7 | -0.67304 | 0.000375 |
| % Perforin$^{+}$ T cell 3 | % CD127$^{+}$ T cell 4 | 0.666957 | 0.000435 |
| % Perforin$^{+}$ T cell 3 | % Perforin$^{+}$ T cell 7 | 0.658261 | 0.00054 |
| % IL-6$^{+}$ Myeloid 2 | % Perforin$^{+}$ T cell 3 | -0.64261 | 0.000779 |
| % CD127$^{+}$ T cell 4 | % Perforin$^{+}$ T cell 7 | 0.642609 | 0.000779 |
| % Perforin$^{+}$ T cell 3 | % CD38$^{+}$ T cell 4 | 0.636522 | 0.00089 |
| % Perforin$^{+}$ T cell 3 | % GranzymeB$^{+}$ T cell 7 | 0.632174 | 0.000972 |
| % GMCSF$^{+}$ Myeloid 2 | % Perforin$^{+}$ T cell 3 | -0.62783 | 0.00106 |
| % CD127$^{+}$ T cell 4 | % GranzymeB$^{+}$ T cell 7 | 0.625217 | 0.001108 |
| % CD38$^{+}$ T cell 4 | % CD127$^{+}$ T cell 4 | 0.61913 | 0.001256 |

### Table 3: Myeloid composition analysis

| **Subset** | **log2FC** | **BH-adjusted p** |
|---|---|---|
| Myeloid1 | 2.46849 | 6.80E-06 |
| Myeloid4 | 2.520732 | 5.97E-05 |
| Myeloid2 | -0.8106 | 0.008 |
| Myeloid5 | -0.57069 | 0.029 |
| Myeloid6 | 0.667078 | 0.082 |
| Myeloid3 | 0.404494 | 0.086 |
